# EssentCell: Discovering Essential Evolutionary Relations in Noisy Single-Cell Data

**DOI:** 10.1101/2025.04.12.648524

**Authors:** Adiesha Liyanage, Robyn Burger, Allison Shi, Braeden Sopp, Binhai Zhu, Brendan Mumey

## Abstract

Single-cell sequencing (SCS) enables investigating tumor evolution at a single cell resolution. A common type of analysis to investigate evolutionary structure from an SCS experiment is to determine a phylogenetic tree structure from the data. This problem has been well-studied under the assumption that mutations only accumulate in the evolution of cancer and there is a simple characterization of when the data is compatible with a perfect phylogeny based on the absence of a special “conflict” submatrix. SCS data can be represented as a binary matrix, where the *ij*-th entry indicates whether cell *i* has mutation *j*. In practice, SCS data is noisy, so a natural question is what is the minimum number of entries to flip in the data matrix, in order that the matrix becomes “conflict-free” and thus compatible with a perfect phylogeny. Furthermore, the false positive rate is orders of magnitude smaller than the false negative rate, so that at most a few false positives occur with high probability. We consider a variation of the minimum-flip problem in which the number of false positives in the solution is a parameter. Often, there can be multiple optimal solutions, so a natural question is what relations are true among all optimal solutions for a small range of possible false positives values; we call such relations *`essential`*. In this work, we propose an efficient algorithm based on integer linear programming to determine all essential relations in the data. We test our software tool, `EssentCell`, on several data sets and discuss the results found.

## 1 Introduction

Cancer is an evolutionary disease. The clonal theory [22] states that most tumors start from a single cell that acquires some growth advantage through genetic mutation. Often, a mutated initial cell proliferates, forming a population of genetically identical clones. These genetic mutations lead to an increase in genetic instability among tumor cells, which increases the likelihood of further genetic mutations and chromosomal abnormalities. As such, growing tumor cells produce more cells with additional genetic changes. Most of these mutated cells die off or are out-competed; however, some tumor cells prevail because of selective advantages (e.g., faster growth, the ability to evade the immune system, etc.), becoming the precursors of new subpopulations within the tumor. This cycle of genetic mutation and selection processes continues throughout the life of the tumor, gradually selecting for increasingly aggressive and malignant tumor cell populations and resulting in a tumor that is comprised of multiple subpopulations with distinct genetic profiles. Thus, accurately predicting the evolutionary history of cancerous cells is highly relevant to learning more about tumor initiation, progression, metastasis, and other related phenomena.

In SCS experiments, a set of single cells is physically taken from a tumor and sequenced. After the mutation calling step, the data can be viewed as a binary mutation matrix, where rows and columns correspond to individual cells and mutations of interest. If mutation *j* is present in cell *i*, then the entry at row *i* and column *j* is equal to 1, otherwise 0. Under the infinite site assumption, if the data has no noise, obtaining an evolutionary history is equivalent to solving the perfect phylogeny problem, for which there is a well-known linear-time algorithm [13]. As previously mentioned, SCS data often contains sequencing errors arising from the amplification cycle. When a mutation goes undetected, the resulting error is classified as a false-negative (FN). Conversely, a false-positive (FP) occurs when a cell is incorrectly identified as having a mutation that it does not possess. In SCS data, the FN error rate can range between 0.1 and 0.3, while the FP rate is generally below 0.001 [20]. The presence of such errors in binary matrices can introduce conflicts, which renders the construction of a perfect phylogeny impossible [3] (also see Def. 3.1 below). This motivates the computational problem to determine the minimum number of entries that must be changed (henceforth called “bit-flips”) for the observed matrix to become conflict-free.

We consider a variation of the minimum-flip problem in which the number of 1 → 0 bit-flips (FPs) is fixed as a parameter *k* and the objective is to minimize the number of 0 → 1 bit-flips (FNs), termed the *MinFlip-k* problem. Often, there can be multiple optimal solutions, each corresponding to a different perfect phylogeny relationship on the cells. A natural question is what relations are true among all such pefect phylogenies, i.e. is cell *v* always a descendant of cell *u*? We call such conserved relations *k-essential* (see Def. 3.3). We assume that *k* can range up to some maximum value *κ*, determined by a standard Chernoff bound probability calculation, based on the given FP error rate of the SCS technology. To illustrate this, consider a mutation matrix that contains 1000 1-entries, with a FP error rate of 0.0005; the probability of finding more than two false positives in the input matrix is at most 0.019 (Chernoff bound). Thus, we discount the events that there are more than 2 false positives, while considering the essential relations that appear in all optimal solutions the problem with *k* ∈ [0, 2]. Those relations that are true in all optimal solutions for all *k* in the range [0*, κ*] are considered [0*, κ*]*-essential* (see Def. 3.4). Our contributions include a fixed-parameter tractable algorithm to determine the *k*-essential relation, as well as a more practical algorithm based on integer linear programming and group-testing. We test our software tool, `EssentCell`, which implements the second algorithm, on several data sets and discuss the results found.

## 2 Related Work

In the past decade, sequencing methods have made rapid advances, leading to the development of computational models to study tumor evolution. Early methods were mainly focused on bulk sequencing data, which is obtained by sequencing a mixture of DNA originating from a large number of cells. This approach has an inherent drawback that it is unable to extract information about the cell of origin of a particular read. For example, this method might output several topologically different evolutionary trees that are able to describe the data equally well, but make it impossible to unambiguously differentiate between them [23].

The alternative group of methods that infer evolutionary history focuses mainly on single-cell sequencing (SCS) data. Due to the fact that SCS data is better at producing high-resolution evolutionary trees, SCS has attracted much attention in tumor phylogenetics [20]. Compared with SCS data, bulk sequencing data is less costly; however, it results in a set of equally likely phylogenies, making it harder to uniquely determine the correct evolutionary tree. Conversely, SCS data, while producing high-resolution evolutionary trees, is subjected to the elevated noise rates present in SCS data sets, doublets (multiple cells are sequences as a single cell), and the high cost of sequencing. Due to these reasons, analyzing SCS data is not straightforward, as we need to develop computational methods that account for noise rates. Moreover, techniques for inferring phylogenies have been developed for both bulk-sequencing, single-cell data [7, 10, 17] and combined joint inference [19, 21, 29].

In [3], the authors considered the combinatorial problem of determining the minimum number of bit-flips needed to convert a binary matrix into a conflict-free matrix and proved the problem to be NP-hard, even if the type of bit flip is restricted to just 0 → 1 or 1 → 0. Recently, this problem has attracted attention again under a slightly modified definition in which the goal is to find the maximum likelihood conflict-free matrix [4,9,10,21,24]. Several approaches have been proposed to solve this version of the problem, employing techniques such as constrained satisfaction programming, mixed-integer linear programming, Markov chain Monte Carlo search, and others. In [2], a deep learning method was used to decide whether the SCS data given indicate a linear or branch evolutionary process. Then, in [28], it was shown that determining the minimum number of 0 → 1 bit flips to obtain a linear perfect phylogeny is NP-complete. Furthermore, they provided a constraint optimization problem formulation to classify an evolutionary trajectory as linear or branched.

Optimization problems often admit multiple optimal solutions. Rather than focusing on a single specific solution, one can ask what properties are present in all optimal solutions. In the literature, these properties are often called by various names, such as “safe” ( [25]), “persistent” ( [16], [5]), or “reliable” ( [26]). The study of safe properties originated in bioinformatics in the 1990s with those amino-acid pairs that are common to all optimal and suboptimal alignments of two protein sequences. In genome assembly, we see this technique applied to “unitigs,” non-branching paths in an assembly graph [18]. Later, in [25], Tomescu and Medvedev formalized “safe” strings that appear in all solutions to a genome assembly problem formulation, expressed as a certain type of walk in a graph. In [8], Dias et al. proposed a method to compute all safe solutions for the NP-hard problem of minimum flow decomposition, applied to RNA sequence assembly.

## 3 Preliminaries

Let *n* be the number of single cells sequenced, and *m* be the number of distinct mutations discovered during the sequencing process. The SCS data is given as a binary mutation matrix *D* ∈{0, 1} *^n×m^* where *D_i,j_* = 1 if cell *i* contains mutation *j*, and 0 otherwise. Next, we formally define a conflict in the binary matrix *D*.

### Definition 3.1.

*A conflict in the mutation matrix D is a triplet of rows i, j, k and a pair of columns p, q with the following pattern:*

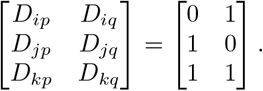

As mentioned previously, the rows of input matrix *D* are leaves of a perfect phylogenetic tree if and only if *D* is conflict-free matrix [14]. Due to the presence of errors, real mutation matrices are unlikely to be conflict-free. Since the false positive error rate is considerably lower compared to the false negative error rate, we define the minimum flip problem under the parameter *k*, where *k* represents the fixed number of 1 →0 bit flips that must be performed while minimizing the number of 0 →1 bit flips. The reasoning behind this choice is that it allows us to consider a range of problem instances with different values of *k* and determine the relations that hold among all optimal solutions. We will use *X* to denote the corrected form of *D*, i.e. a corrected data matrix that is conflict-free.

### Definition 3.2.

*We define the* minimum flips with exactly *k* 1-flips problem *(MinFlip-k) to determine a* {0, 1}*^n×m^* conflict-free matrix X such that X is obtained from D by performing a minimum number of 0 → 1 *flips, while requiring exactly k* 1 → 0 *flips.*

### 3.1 The Essential Partial Order

We are interested in determining the evolutionary relationships among the set of perfect phylogenies that are associated with each optimal solution to MinFlip-*k*. With this in mind, we define the *k-essential* order relation 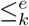 on the set of cells as follows:

#### Definition 3.3.

*Let* (*D, k*) *be an instance of MinFlip-k. We say 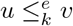 if and only if X_uj_* ≤ *X_vj_ for all columns j in all optimal solutions X.*

#### Property 3.1.

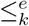 *is transitive.*

*Proof.* This can be easily seen from the definition: if 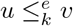 and 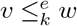,then clearly 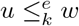 since ≤ on each column position is transitive.

**Property 3.2. 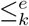** *is reflexive.*

*Proof.* Consider any cell *u*. For each column *j*, clearly *X_uj_* = *X_uj_*, hence 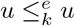.

Let *κ* be the maximum number of FPs considered. We define the [0*, κ*]*-essential* order relation 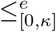 on the set of cells as follows:

#### Definition 3.4.

*We 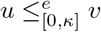 if and only if 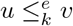 for all k* ∈ [0*, κ*].

It is possible that two distinct cells *u* and *v* can be *strongly-connected*, i.e. 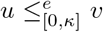 and 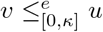.

For this reason, we will collapse the cell space into *strongly-connected components* (SCCs) under the relation 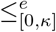.

**Property 3.3. 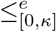** *on SCCs is a partial order.*

*Proof.* By definition, 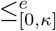 on SCCs is anti-symmetric, and is easily seen to be reflexive and transitive.

### 3.2 An FPT algorithm to compute 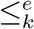

In [3], Chen et al. gave an *O*^*^(6*^t^*) time *fixed-parameter tractable* (FPT) algorithm for the minimum flip problem, where both 0→ 1 and 1→ 0 flips are allowed and *t* is the minimum number of flips; if only 0→ 1 flips are permitted, the running time drops to *O*^*^(2*^t^*) time.^1^ Their algorithm is based on an equivalent formulation on a bipartite graph: given *D* we convert it into a bipartite graph *G* = (*A, B, E*), where *A* contains all the rows (cells) and *B* contains all the columns (mutations); moreover, there is an edge (*i, p*) ∈ *E*, for *i* ∈ *A* and *p* ∈ *B*, if and only if *D_i,p_* = 1. Then a forbidden structure (called an ℳ-path) in *G* is a cycle-free 4-path ⟨*i, p, j, q, k*⟩ where {*i, j, k A}* and {*p, q}* ∈ *B*. To make *D* conflict-free we need to destroy all forbidden ℳ-paths. Specifically, to eliminate ⟨*i, p, j, q, k*⟩, we can either insert one of the two edges (*i, q*) and (*k, p*), or delete one of the 4 edges in the 4-path. The algorithm proceeds by picking any ℳ-path in *G* and making one of the 6 choices available to eliminate it. The search space is thus a tree with a branching factor of 6; if breadth-first search is used, then all optimal solutions will be found at depth *t* in the tree.

We can adapt this algorithm to find the *k*-essential relation as follows: In our case we must enumerate all solutions to MinFlip-*k*. This requires running the Chen et al. algorithm (running in *O*^*^(2*^t^*) time, as only 0→ 1 flips are allowed) for all possible ways (at most 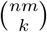) to choose *k* false positives. Some choices might be infeasible (i.e. 0→ 1 flips cannot undo 1 c→0 flips; so some conflicts may not be removable); we can detect these and not run the algorithm for those cases. For the false positive choices that remain, we can run the searches in parallel to determine all optimal solutions with the minimum number (*t*) of 0 → 1 flips. We have,

#### Theorem 3.1.

*MinFlip-k can be solved in O*^*^((*nm*)*^k^*2*^t^*) *time, where t is minimum number of* 0 →1 *flips required.*

Note that if we consider *k* as a fixed constant (rather than a parameter) and set *t* as a parameter, then this corollary also implies that MinFlip-*k* is FPT. The relation 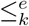 is found by intersecting the row-orders of all optimal solutions (in *O*(*n*^2^) time per solution found), so we also have,

#### Corollary 3.1.

*The k-essential relation for a single cell data set D can be found in O*^*^((*nm*)*^k^*2*^t^*) *time, where t is minimum number of* 0 → 1 *flips required.*

However, when *k* is not a very small constant or *D* is somewhat noisy (*t* large), this algorithm is also impractical. Therefore, in the following, we propose a different algorithm based on integer linear programming.

### 3.3 ILP formulation for MinFlip-*k*

We first provide an integer-linear program (ILP) for MinFlip-*k* based on the ILP given in [15] for the minimum flip problem. The variables for the ILP include the predicted conflict-free matrix *X* ∈ {0, 1} *^n×m^* as well as some auxiliary variables *B_p,q,a,b_* ∈{0, 1}, where 1 ≤ *p < q* ≤ *m* and (*a, b*) ∈ {(0, 1), (1, 0), (1, 1)}. We add constraints to ILP such that *B_p,q,a,b_* = 1 if and only if there exists a row *i* such that *X_ip_* = *a* and *X_iq_* = *b*. The full constraints to the ILP are given below:

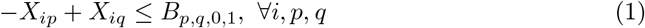

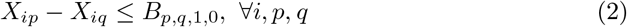

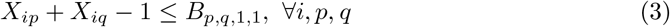

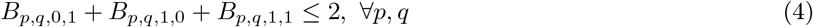

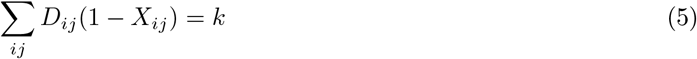

Constraints (1) through (4) enforce that *X* is conflict-free, while (5) ensures that the number of 1 → 0 flips is exactly *k*. Finally, the objective function is

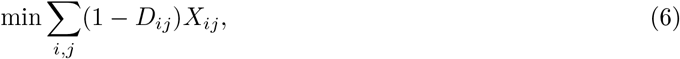

which minimizes the total number of the 0 → 1 flips.

## 4 A group-testing algorithm to compute 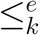

We next describe an algorithm to compute the *k*-essential partial order based on a *group-testing* approach using the ILP from Section 3.3. We first describe an approach to test if a single edge (*u, v*) belongs to the relation; that is for two given cells *u* and *v*, we would like to determine whether 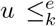 *v*, i.e. we would like to know if all optimal solutions to the MinFlip-*k* instance require that *X_uj_* ≤ *X_vj_* for all columns *j*. We will check this using the contrapositive version of the statement, namely that there are no optimal solutions *X* such that *X_uj_ > X_vj_* for some column *j*. We can test this contrapositive statement by modifying the ILP from Section 3.3; we introduce additional {0, 1}-variables {*z_j_*} and the following constraints:

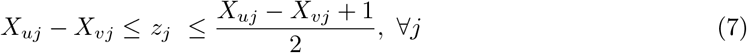

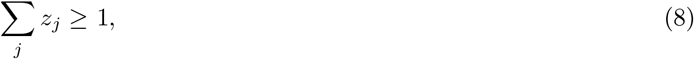

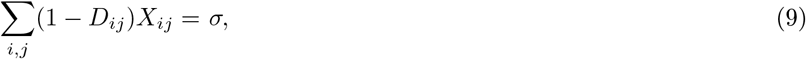

where *σ* is the optimal objective value for the original ILP. Constraint (7) ensures that *z_j_* = 1 if and only if *X_uj_ > X_vj_*, while (8) requires that some *z_j_* = 1. Let `TestEss`(*k, u, v*) be this modified ILP. We have:

**Lemma 4.1. 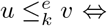** *TestEss*(*k, u, v*) *is infeasible.*

*Proof.* The proof follows directly from the fact that checking 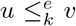 is equivalent to checking the contrapositive version of Definition 3.3, which, in turn, is true if and only there are no feasible solutions *X* to the original ILP with *X_uj_ > X_vj_* for some *j* that achieve the objective value *σ*.

In principle, we could determine 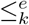 by testing every pair of cells using Lemma 4.1, or by using a more sophisticated approach such as [6] that has *O*(*wn* log *n*) query complexity and *O*(*w*^2^*n* log *n*) time complexity, where *w* is the *width*^2^ of the relation. In this work, we chose to use a *group-testing* approach based on a similar ILP test: Suppose *u* is a cell and let 𝒱 be a set of additional cells not containing *u*. We modify the ILP from Section 3.3 in a similar way to above; for each *v* ∈ *𝒱*, we create a new variable 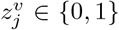 that must satisfy the constraints (7) and (8). Let `TestEss`(*k, u, 𝒱*) be this modified ILP. We have:

### Lemma 4.2.

*TestEss*(*k, u,* 𝒱) *is feasible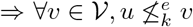.*

*Proof.* We prove the contrapositive: 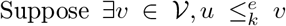 and `TestEss`(*k, u,* 𝒱) is feasible. This contradicts Lemma 4.1, since `TestEss`(*k, u,* 𝒱) contains all the same constraints that made `TestEss`(*k, u, v*) infeasible.

Lemma 4.2 suggests a recursive strategy to determine 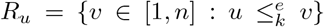:Initially set 𝒱 = [1*, n*] and test whether 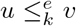 for some *v* ∈ 𝒱 (we know *u* ∈ *R_u_* by Property 3.2). If so, partition 𝒱 into two subsets of equal size and test each subset recursively. Any subset that does not contain any elements from *R_u_* will be eliminated with one ILP test. When |𝒱 | = 1, if `TestEss`(*k, u, v*) is infeasible, we learn 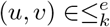.By Property 3.1 (transitivity), we can also add `in`(*u*) × `out`(*v*) to 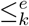 (we define `In` 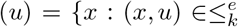,and out 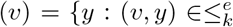). Algorithm 1 implements this strategy and the general flow of the strategy is illustrated in Figure 1. Adding these transitive edges may also imply that some *v* ∈ V are already reachable from *u* in a subsequent `TestEss`(*k, u,* 𝒱) call; line 2 in Algorithm 1 removes any such *v* where (*u, v*) is already a known relation edge. The complete procedure to find 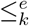 is given in Algorithm 2.

**Figure 1.**
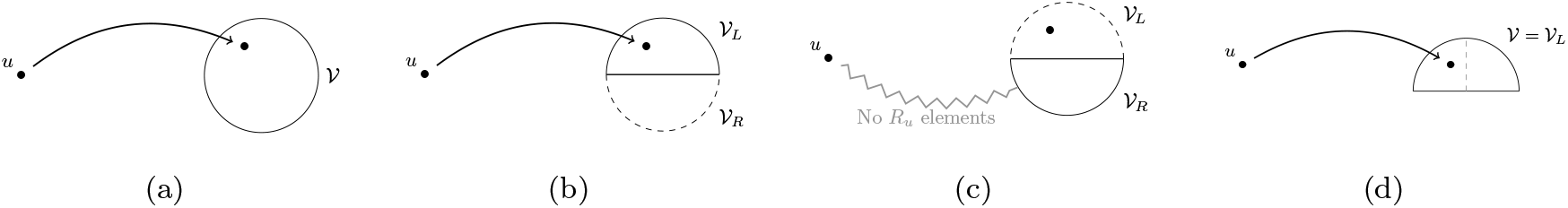
The strategy suggested by Lemma 4.2. We start with 𝒱 as depicted in (a). We divide 𝒱 into roughly equal sized partitions 𝒱*_L_* and 𝒱*_R_* then test each separately as in (b) and (c). Any partition with an element in *R_u_* is split again as indicated by the gray dashed line in (d).

We run Algorithm 2 for each *k* ∈ [0*, κ*], and thus determine 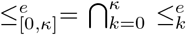. Finally, we find the strongly-connected components of 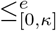 (collapsing each to a single vertex) and then take the *transitive reduction* to more compactly represent the relation. The transitive reduction removes edges that can be inferred by transitivity and is unique for directed acyclic graphs [1].

### Algorithm 1

Group-testing algorithm.

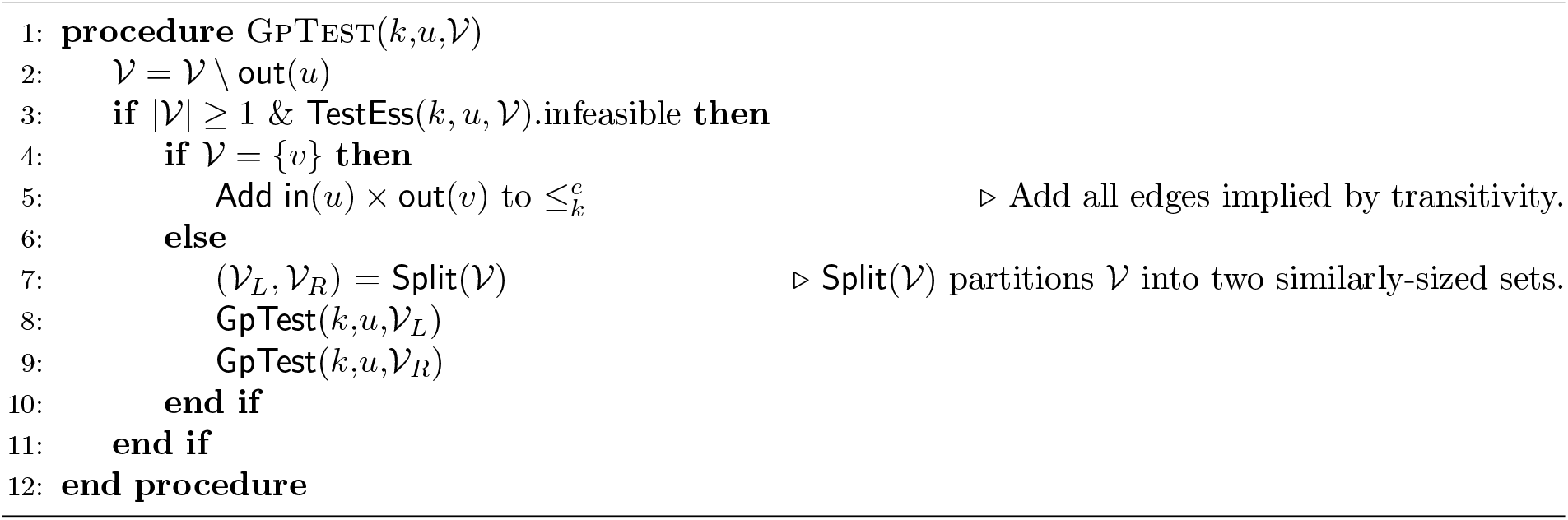

### Algorithm 2

Full procedure to find 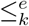.

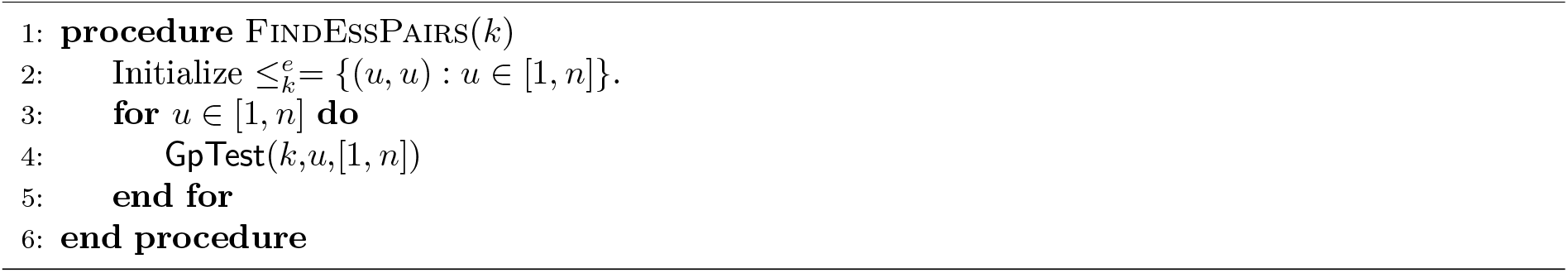

## 5 Experimental Results

We tested our tool, `EssentCell` — which is publicly available at https://github.com/msu-alglab/essentcell —, on three data sets (see Table 1). Computational efforts were performed on a local HPC cluster ^3^; each experiment was run on a node with an Epyc Genoa/Rome X86 CPU with 64 Gb of memory allocated. We implemented the EssentCell tool in Python using Gurobi Optimizer [12] as the ILP solver. Two data sets (Patient 2 and Patient 6) were taken from a set of six acute lymphoblastic leukemia (ALL) patients in [11]; these were chosen as test cases for the tool Phyolin [28] to investigate that tool’s predictions of linear versus branched evolutionary structure. A third dataset (ER-tumor) was taken from a *ER*^+^ breast cancer sample in [27]. Note that we apply a timeout to ILP calls for groups larger than 2, while allowing the optimizer to fully solve ILPs for groups of size 1. If a timeout occurs, the algorithm splits the group into smaller subgroups and continues testing, ensuring that the optimal solution is still found.

**Table 1:** Data Sets tested: *n* is the number of cells, *m* is the number of mutations, *k* is number of false positives. A Chernoff bound is used to compute the probability of more than *k* false positives; the selected values for *κ* are shown in bold. The number of ILP calls made by Algorithm 2 is given, as well as the percent of ILP calls saved in comparison to the naive approach without group testing. Lastly, the average time per call is given.

| Data Set | $n$ | $m$ | $k$ | $\Pr(\#FP > k)$ | ILP calls | ILP calls saved (%) | Avg. time per ILP call |
| --- | --- | --- | --- | --- | --- | --- | --- |
| Patient 6 | 146 | 10 | 0 | 0.404 | 5932 | 44.0% | 10.4 s |
|  |  |  | 1 | 0.055 | 2451 | 76.8% | 11.6 s |
|  |  |  | **2** | **0.004** | 2784 | 73.7% | 15.6 s |
| Patient 2 | 115 | 16 | 0 | 0.310 | 2863 | 56.3% | 18.4 s |
|  |  |  | 1 | 0.027 | 2840 | 56.6% | 21.3 s |
|  |  |  | **2** | **0.001** | 2914 | 55.6% | 28.2 s |
| ER-tumor | 47 | 41 | 0 | 0.314 | 685 | 36.6% | 66.8 s |
|  |  |  | 1 | 0.028 | 661 | 38.9% | 276.9 s |
|  |  |  | **2** | **0.002** | 661 | 38.9% | 433.4 s |

To determine the appropriate *κ* value for each data set, we used a Chernoff bound to estimate the probability that the number of false positives (#FP) exceeded *k* for the values shown in Table 2. This calculation uses the number of 1 s in each matrix *D* and assumes an independent false positive probability of Pr(1 → 0) = 0.0005. We choose *κ* to obtain a less than 1% chance that more than *κ* false positives occurred in *D* (chosen values are bolded in Table 2). Next, we ran Algorithm 2 to determine the *k*-essential related pairs for each data set, and recorded the total number of ILP calls made (the initial ILP call to find the objective value plus all calls made to test the feasibility of the `TestEss` ILP). The strongly connected components of 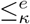 were computed and collapsed to be single vertices, and the *transitive reduction* of the resulting graph was found, resulting in the graphs shown in Figure 2. Finally, the *width* of each resulting graph was computed.

**Table 2:** Summary of the essential relations found for the *κ* values selected from Table 1. The Phyolin [28] prediction, poset width and number of nodes are given.

| Data Set | Phyolin prediction | Width | Nodes |
| --- | --- | --- | --- |
| Patient 6 | Linear | 4 | 14 |
| Patient 2 | Branched | 3 | 20 |
| ER-tumor | Branched | 13 | 27 |

**Figure 2.**
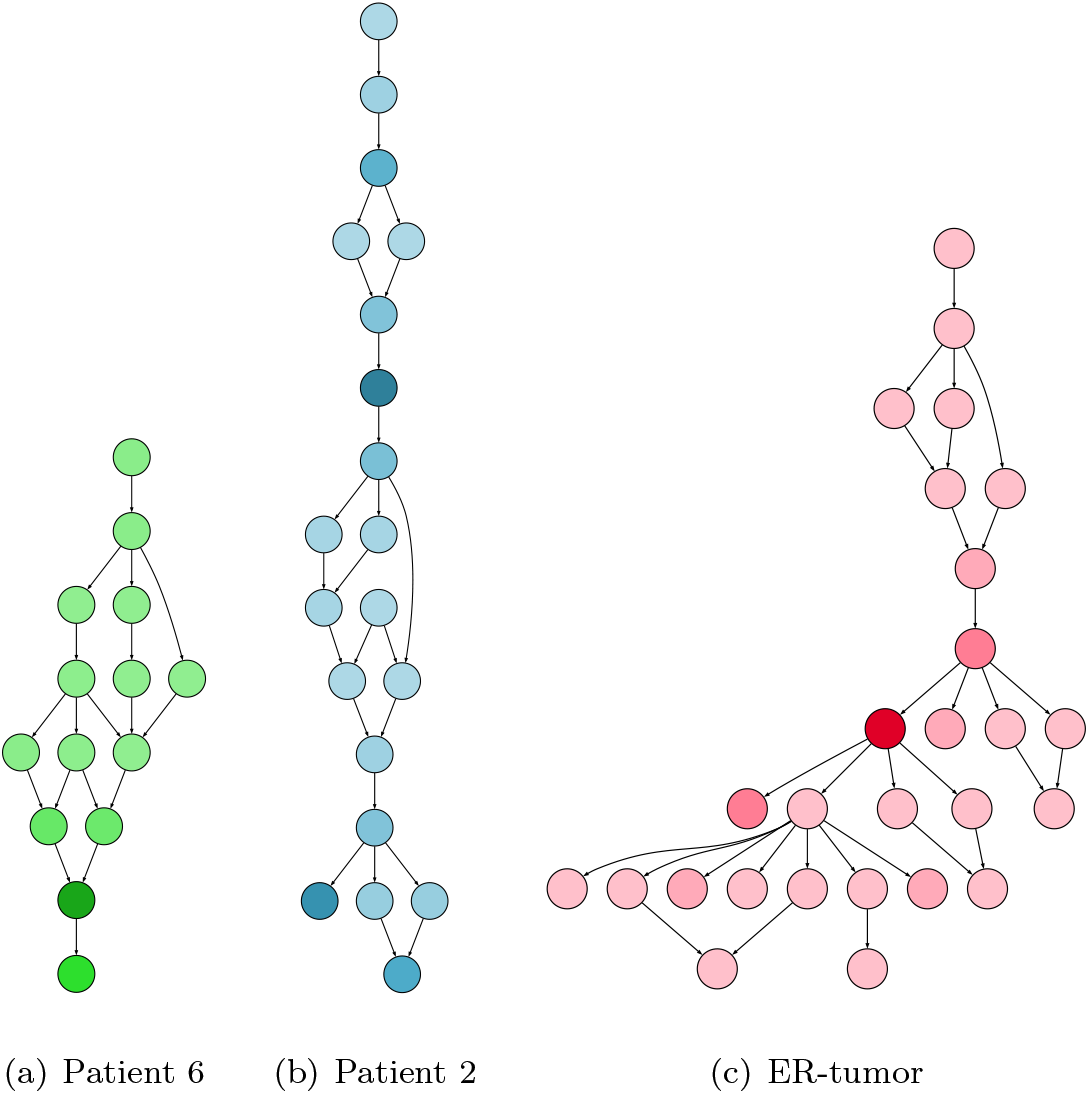
Hasse diagrams of the [0, 2]-essential relation for Patient 6, Patient 2, and ER-tumor data sets. Nodes are shaded according to the number of cells contained. (A Hasse diagram depicts the partial order so that all edges point down; thus the arrowheads are not shown.)

At a high level, the essential relations found have some agreement with the coarser categorical descriptions of Phyolin, e.g., Figure 2(b) has multiple leaf nodes that contain larger number of cells; this structure presumably requires more flips to reach a linear structure than, Figure 2(a) which has a unique leaf.

We also examined the evolution of two specific mutations in the ER-tumor data set, **MARCH11** and **CABP2**. Figure 3 illustrate how these mutations evolve in the [0, 2]-essential relation. A node is colored white if none of cells associated with that node have the mutation. Nodes are black if all node cells are predicted to have the mutation, and at least one had the mutation in the original data matrix (a true positive). Finally, nodes are grey if all node cells are predicted to have the mutation, but none had it in the original data (thus all node cells were false negatives). We selected these two mutations because they were identified as subclonal mutations in [27] that emerged later in tumor evolution. Moreover, cells corresponding to white nodes in Fig. 3 contain more clonal mutations and fewer subclonal mutations, whereas black and gray nodes contain clonal mutations along with some subclonal mutations. This suggests that these cells may have acquired the subclonal mutations at a later stage in tumor evolution. Moreover, [27] notes that the **MARCH11** mutation was present in a larger subset of the sequenced cells, whereas **CABP2** was detected in fewer sequenced cells, suggesting that **CABP2** may have emerged even later in tumor evolution. Our findings agree with this, as seen in Figure 3.

**Figure 3.**
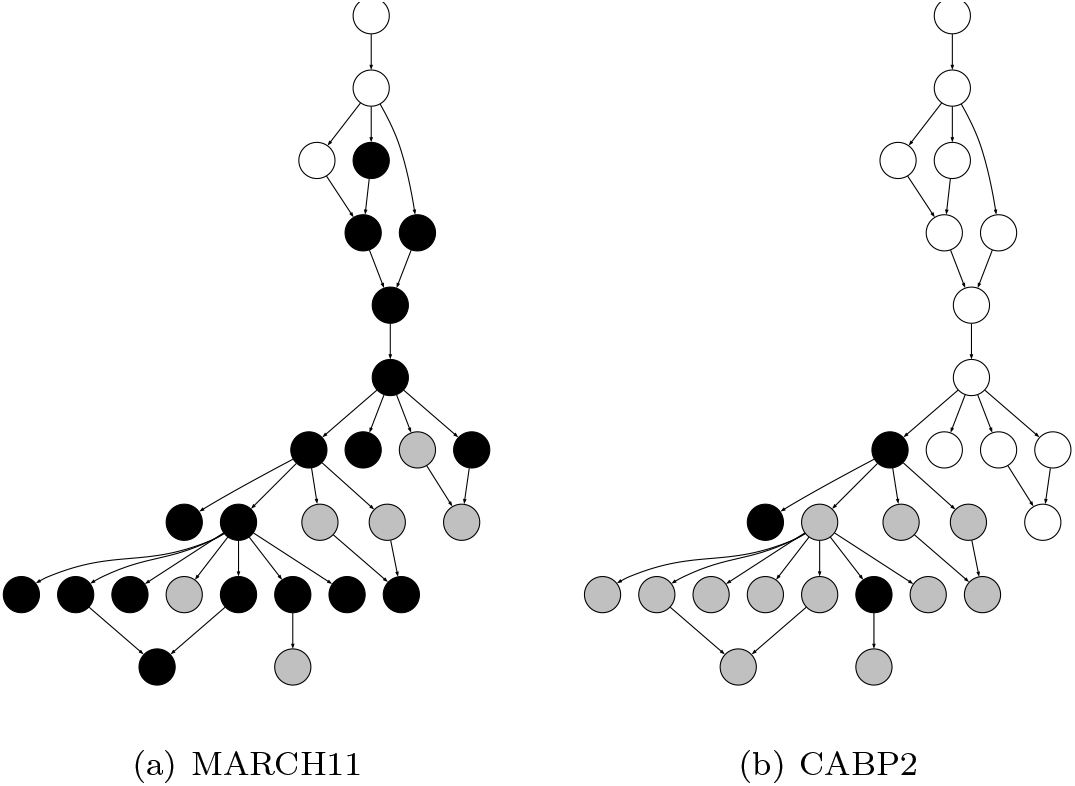
Hasse diagrams of the [0, 2]-essential relation for the ER-tumor data set, illustrating the presence of two mutations: MARCH11 and CABP2. White nodes indicate the cells in the given node do not have the mutation (no optimal solutions predict this). Black nodes indicate that the mutation is present in the data for some cells associated with a given node, and predicted for all cells in the node, whereas gray nodes indicate the mutation is predicted for all cells in the node, but not present in the original data (thus such cells were false negatives for the mutation).

## 6 Discussion

Single cell data is much more likely to have false negatives versus false positives; this motivated us to consider what relations hold in all optimal solutions when we want to minimize the number of false negatives, while allowing a small number of false positives. We framed this question as computing an *essential relation* that induces a partial order on the cells and helps elucidate the evolutionary structure that is consistent with all optimal solutions. Interestingly, these essential relations do not always form a tree, unlike the assumed underlying phylogenetic relation between the cells; this is due to the fact that there can be multiple optimal phylogenetic relations, and the essential relation captures those relations that hold among all possible optimal trees. Besides investigating these relations for new biological insights, it is interesting to consider how to further optimize the algorithms for finding the essential order, where the goal is to minimize the number of comparisons or group-tests. The perfect phylogeny assumption idealizes the problem as real data may include recurrent or back mutations and some tumors may exhibit copy number alterations (CNAs) that can create loss of variations through loss of heterozygosity. The essential relation idea could be applied to other models of tumor evolution for which an optimization problem is formulated and an efficient test for “essentialness” is available, in at least a pairwise formulation.

## Acknowledgements

This work has received funding from NSF awards 2243010 (REU site), 2309902, and 2307572.

## Footnotes

1 The *O*^*^ notation suppresses the polynomial factor related to the input size.

2 The width of a directed graph *G* is defined as the maximum size of an antichain in *G* and is equal to the size of a minimum path cover of *G*, by Dilworth’s theorem.

3 Tempest High Performance Computing System, operated and supported by University Information Technology Research Cyberinfrastructure at Montana State University.

